# Surveillance of Shiga toxin-producing *Escherichia coli* associated bloody diarrhea in Argentina

**DOI:** 10.1101/2021.01.07.425824

**Authors:** Marta Rivas, Mariana Pichel, Mariana Colonna, Adrián López Casanello, Laura F. Alconcher, Jimena Galavotti, Iliana Principi, Sofía Pérez Araujo, Flavia B. Ramírez, Gladys González, Luis A. Pianciola, Melina Mazzeo, Ángela Suarez, Sebastián Oderiz, Lidia F. R. Ghezzi, Diego J. Arrigo, José H. Paladini, María R. Baroni, Susana Pérez, Ana Tamborini, Isabel Chinen, Elizabeth S. Miliwebsky, Fernando Goldbaum, Luciana Muñoz, Linus Spatz, Santiago Sanguineti, the EPI-HUS Investigation Team

## Abstract

In Argentina, the hemolytic uremic syndrome associated with Shiga toxin-producing *Escherichia coli* (STEC-HUS) infection is endemic, and reliable data about prevalence and risk factors are available since 2000. However, information about STEC-associated bloody diarrhea (BD) cases is limited. A prospective study was carried out in seven tertiary-hospitals and 18 Referral Units from different regions, aiming to determine (i) STEC-positive BD cases frequency in 714 children aged 1 to 9 years old; and (ii) rate of progression to HUS. The number and regional distribution of STEC-HUS cases assisted in the same hospitals and period was also assessed. A total of 29 (4.1%) STEC-positive BD cases were confirmed by Shiga Toxin Quik Chek (STQC) and/or mPCR. The highest frequencies were found in the Southern region (Neuquén, 8.7%; Bahía Blanca, 7.9%), in children between 12 and 23 month of age (8.8%), during summertime. Four (13.8%) cases progressed to HUS, three to five days after BD onset. Twenty-seven STEC-HUS children mainly under 5 years old (77.8%) were enrolled, 51.9% were female; 44% were Stx-positive by STQC and all by mPCR. The most common serotypes were O157:H7 and O145:H28 and prevalent genotypes were *stx*_2a_-only or associated, both among BD and HUS cases. Considering the endemic behavior of HUS and its impact on public health, it is important to have updated information about the epidemiology of the diarrheal disease for early recognition of infected patients and initiation of supportive treatment. Finally, it also gives the opportunity to respond to outbreak situations effectively and in timely manner.

## INTRODUCTION

Shiga toxin-producing *Escherichia coli* (STEC) are a heterogeneous group of foodborne pathogens and an important cause of morbidity and mortality, with associated loss of life years and diminished health-related quality of life. The clinical manifestations of infection range from symptom-free carriage to nonbloody and bloody diarrhea (BD), hemorrhagic colitis, and hemolytic-uremic syndrome (HUS) (1). HUS is a systemic thrombotic microangiopathy, involving acute kidney failure that may cause death or end-stage renal disease, a serious chronic condition that reduces life expectancy (2, 3).

The linkage between STEC infection and the development of HUS was established in early 1980s (4, 5). Shiga toxin 1, and/or Shiga toxin 2 have been demonstrated to be the primary virulence traits responsible for human disease (6) and many subtypes have been described (Stx1a, Stx1c, Stx1d, Stx2a, Stx2b, Stx2c, Stx2d, Stx2e, Stx2f, Stx2g, and the recently described Stx2h, Stx2i, Stx2k) (6, 7). Strains producing Stx2a, Stx2c, or Stx2d have been reported to be more pathogenic than those strains producing Stx1 subtypes alone or both Stx1 and Stx2 (8). In addition, a mosaic of different virulence traits, comprising several adhesins and other toxins that may play a role in pathogenesis, has also been described (9).

STEC isolates belong to a large number of O:H serotypes, and O157:H7 is the most prevalent serotype associated with severe human diseases. However, other non-O157 serogroups as O26, O45, O103, O111, O121, and O145 have been identified (10, 11).

After STEC infection, the mean incubation period is 3 days (ranging from 2 to 12 days) (12, 13). The predominant symptom is diarrhea, being bloody in 80% of the patients. Abdominal pain is more severe than in other cases of bacterial gastroenteritis and most patients infected with O157:H7 do not present fever. Nausea and vomiting are sometimes reported. (14). Diarrhea usually improves in one week, however, approximately 15% of cases evolve to HUS (2 to 14% in sporadic cases, up to 20% during outbreaks) (15). STEC-HUS represents approximately 90% of the total HUS cases in children.

In Argentina, STEC-HUS is endemic, with approximately 350 new cases reported annually. In the last 10 years, the incidence has ranged from 8 to 10 cases per 100,000 children under 5 years of age and lethality was between 1 and 4%. During 2018, 339 HUS cases were notified, with a rate of 0.74 cases per 100,000 inhabitants. The distribution of cases showed a marked difference between the different regions of the country. (Boletín Integrado de Vigilancia N° 463–SE 34/2019. Available online: https://www.argentina.gob.ar/sites/default/files/biv_463_cuatrisemanal.pdf (accessed on 10 June 2020).

At present and given the requirement of mandatory notification of HUS since 2000, there are reliable data about the prevalence and risk factors of this disease in our country (16). However, the available data on the frequency of STEC-associated BD cases, their distribution and the number that evolves to HUS in pediatric patients are limited.

The present study aimed to determine: (i) STEC-positive frequency in BD cases in children between 1 and 9 years old; and (ii) rate of progression to HUS in STEC positive-BD cases within 28 days from the diagnostic confirmation. Additionally, the number and regional distribution of STEC-HUS cases in children aged 1 to 9 years old assisted in the same hospitals during the same period was evaluated.

## MATERIALS AND METHODS

### Study Design

An observational, prospective, descriptive and multicenter surveillance study of pediatric patients with acute BD (first cohort) was conducted between October 10, 2018 and June 30, 2019 at seven participating tertiary-care hospitals (named Reference Sites) and their 18 hospitals of less complexity (defined as Referral Units), located in Buenos Aires, La Pampa, Mendoza, Neuquén, Río Negro and Santa Fe Provinces and Buenos Aires City, Argentina. Screening criteria included children of either gender between 1 and 9 years old who presented acute BD, defined as an increase in the frequency of bowel movements and a decrease in consistency, with the presence of visible blood or blood observable only under a microscope, with less than 2 weeks-duration.

A second cohort of HUS cases aged between 1 to 9 years, with documented history of STEC-positive BD assisted at the same hospitals during the same period was studied. HUS was defined as the triad: microangiopathic hemolytic anemia (fragmented red blood cells in peripheral blood smear - schistocytes, of non-immune origin [negative direct Coombs test]), thrombocytopenia (platelet count < 150,000 cells/mm3), and renal dysfunction (serum creatinine concentration >2 standard deviations above the upper limit of normal for age and/or proteinuria and/or hematuria).

The demographic, clinical and microbiological data of all cases incorporated were obtained from the medical history and related medical records, whether of paper or electronic origin, according to the standards of the Reference Site. A study-specific and validated electronic Case Report Form (eCRF) was used as the only tool for collection, storage and reporting of data. Data recorded during admission were age (in months), gender, origin (province/locality), date of onset of bloody diarrhea, date of collection of the stool sample (identified by a code number for microbiological analysis), and history of antibiotic exposure. Progression to HUS within 28 days from the STEC diagnostic confirmation was evaluated and the date of the diagnosis was recorded.

The study was carried out in compliance with the legal and regulatory requirements, as well as scientific purpose, value and rigor, and the principles established in the Declaration of Helsinki (17) and in the Guide for Good Clinical Practices (ICH Harmonised Tripartite Guideline: Guideline for Good Clinical Practice E6(R2). November 9th, 2016. URL: http://www.ich.org/fileadmin/Public_Web_Site/ICH_Products/Guidelines/Efficacy/E6/E6_R2__Step_4_2016_1109.pdf).

The study protocol was approved by the Institutional Review Boards or the local Ethics Committees as required by local regulation [Disposición ANMAT N° 6677/10. Nov 2010. URL: http://www.anmat.gov.ar/comunicados/dispo_6677-10.pdf] and institutional Standard Operating Procedures. Thus, each Reference Site joined the study as soon as this approval was reached, starting by Hospital Penna of Bahia Blanca, Buenos Aires, in October 2018 and finishing with Hospital Molas of Santa Rosa, La Pampa, in February 2019.

### Laboratory Methods

Dates of reception and processing of the stool sample in the local laboratory, type of sample (fresh stool or rectal swab) and result of the microbiological tests were recorded.

All fecal samples were screened with the SHIGA TOXIN QUIK CHEK (STQC) test (Abbott™, TechLab^®^, Blacksburg, VA) according to the manufacturer’s instructions and the standard guidelines established by the National Reference Laboratory of Argentina, using 2 test procedures: (i) directly on feces (direct method), and (ii) after overnight (16 to 20 h) stationary incubation at 37°C in 8 ml of Gram-Negative (GN) Broth-Hajna (Becton Dickinson, Sparks, MD) (enrichment method). A total of 413/714 fecal samples were processed by an in-house validated multiplex PCR, according to the laboratory capacity (18). These samples were plated onto sorbitol-MacConkey agar (Becton Dickinson) directly, and after enrichment at 37°C for 4 h in trypticase soy broth (TSB) (Becton Dickinson), and TSB supplemented with cefixime (50 ng/mL) and potassium tellurite (25 mg/mL) (CT-TSB). The confluent growth zone and colonies were screened for *stx*_1_, *stx*_2_, and *rfb*_O157_ genes. Isolates with *stx*_1_ and/or *stx*_2_ genes were confirmed as *E. coli* by standard biochemical tests.

STEC infection in HUS patients was established by at least one of following laboratory criteria: (i) screening of Stx1 and/or Stx2 by STQC; (ii) screening of *stx* genes by mPCR and/or STEC isolation; (iii) detection of free fecal Shiga toxin (FFStx); and (iv) detection of anti-O serogroup-specific antibodies O157, O145, O121.

Fecal samples, and *stx*-positive isolates and sera were sent to National Reference Laboratory for confirmation and further characterization (ANLIS/INEI/MP-ARG;2019 http://sgc.anlis.gob.ar/handle/123456789/1682, 19, 20).

### Statistical Analysis

Descriptive and exploratory analysis was performed using R software version 3.5.0 (R Foundation for Statistical Computing, Vienna, Austria). Continuous variables were summarized by means of mean/standard deviation and median/interquartile range (IQR), whereas frequency and proportions were applied to summarize qualitative variables. Categorical variables were compared using the Pearson chi-square test. All tests were two-sided and *P*-values less than 0.05 were considered as statistically significant.

The following proportions and intervals were estimated: 1) proportion of STEC-positive BD by any method (STQC, direct and/or enriched culture, and mPCR) globally and by Reference Site; 2) cumulative incidence of STEC-positive BD evolving to HUS; 3) interval between the onset of BD and seeking medical attention; 4) interval between the onset of BD symptoms and sample collection; 5) interval between sample collection and processing; and 6) frequency of confirmed HUS cases. The performance of the STQC in comparison to the mPCR was also evaluated.

## RESULTS

### Study population

A total of 714 children with BD were included in the study. The median age was 42 months (IQR 24-63; range, 11-119 months), 44.8% (n=320) were female and 4.2% (n=30) received antimicrobial treatment before admission to the hospital. The median interval between the BD onset and seeking medical attention was 0 days (IQR 0-1 days; range, 0-17 days).

Twenty-seven STEC-HUS cases were enrolled. The median age was 37 months (IQR 25-59 months; range 16-104 months) and 51.9% (n=14) were female. For this cohort, the median interval between the BD onset and the STEC-HUS diagnosis was 4 days (IQR 1-6 days; range, 0-10 days).

### Distribution of bloody diarrhea cases by Reference Site

Among the 714 patients, Hospital “Dr. José Penna” of Bahía Blanca enrolled 12.4% of the cases, Hospital “Dr. Humberto J. Notti” of Mendoza 18.8%, Hospital “Dr. Castro Rendón” of Neuquén, 16.1%, Hospital “Sor María Ludovica” of La Plata 21.9%, Hospital Italiano of Buenos Aires City 11.6%, Hospital “Dr. Orlando Alassia” of Santa Fe 11.2%, Hospital “Dr. Lucio Molas” of Santa Rosa, La Pampa 8.0%. Of the total, 61.5% patients were assisted at the Reference Sites and 38.5% at the corresponding Referral Units. The geographic distribution of the BD cases is showed in Fig. 1.

**FIG 1.**
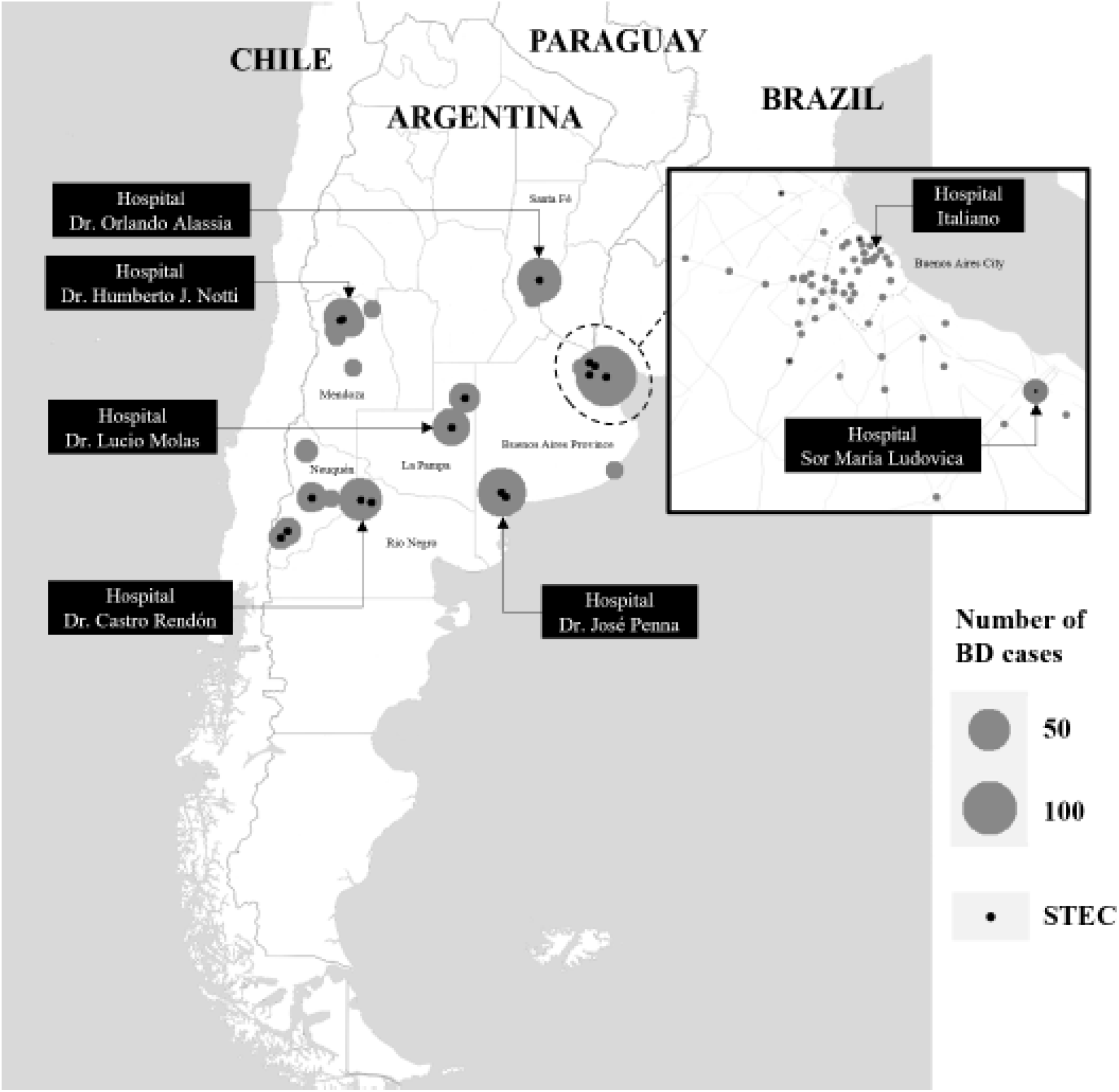
Geographic distribution of bloody diarrhea and STEC-positive bloody diarrhea cases. Map data obtained from © OpenStreetMap contributors. Data is available under the Open Database License https://www.openstreetmap.org/copyright.

### Microbiological and demographic findings

Two hundred and eighty-nine (40.5%) of stool samples were collected on the same day of BD onset, 250 (35%) one day later, while the remaining 175 (24.5%) were collected two or more days after BD onset; 593 (83.1%) were processed by STQC within the first 24 h of sampling.

A total of 29 cases were confirmed by STQC and/or mPCR, resulting in a cumulative incidence of 4.1% (95% CI: 2.7% - 5.8%). Of the 29 STEC-positive BD cases, 21 (72.4%) were detected by the STQC test, 13 (44.8%) by the direct method and other 8 (27.6%) by the enrichment method. By mPCR, 28 cases were *stx*-positive. In the remaining case that tested positive by STQC, the mPCR was not done, however FFStx was detected at the National Reference Laboratory.

Regarding the age group distribution, the highest proportion of STEC-positive BD (8.8%, 14/160) was found in the children 12 to 23 months and significant differences were observed among age groups (chi-square test for equal proportions (α=0.05), *P*=0.0026; pairwise comparisons: 12-23 *vs*. 24-59, *P*=0.0012; 12-23 *vs*. 60-120, *P*=0.0214; 24-59 *vs*. 60-120, *P*=0.5365) (Table 2).

**TABLE 1.**
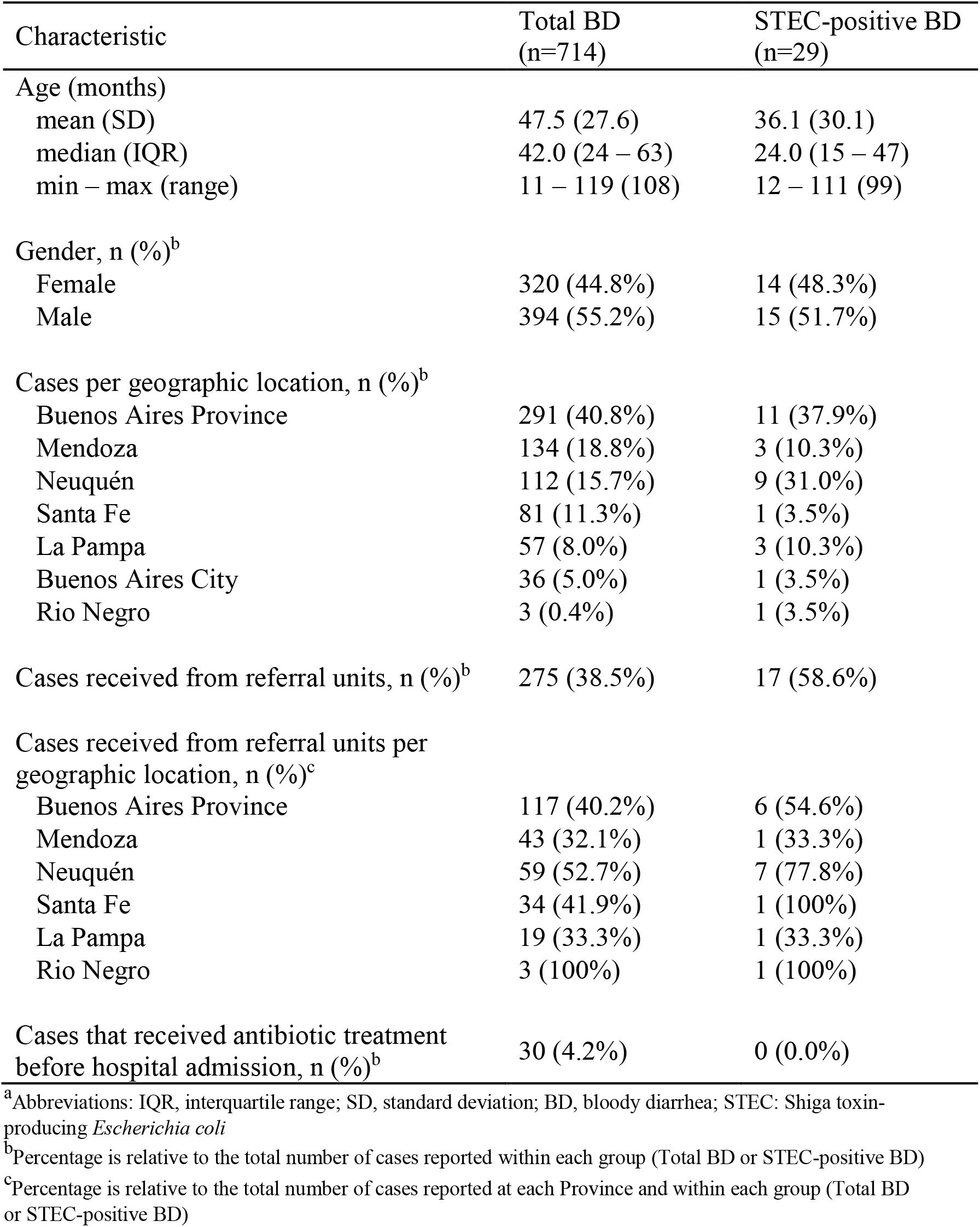
Demographic characteristics of BD and STEC-positive BD cases^a^

| Characteristic | Total BD<br>(n=714) | STEC-positive BD<br>(n=29) |
| --- | --- | --- |
| Age (months) |  |  |
| mean (SD) | 47.5 (27.6) | 36.1 (30.1) |
| median (IQR) | 42.0 (24 – 63) | 24.0 (15 – 47) |
| min – max (range) | 11 – 119 (108) | 12 – 111 (99) |
| Gender, n (%)<sup>b</sup> |  |  |
| Female | 320 (44.8%) | 14 (48.3%) |
| Male | 394 (55.2%) | 15 (51.7%) |
| Cases per geographic location, n (%)<sup>b</sup> |  |  |
| Buenos Aires Province | 291 (40.8%) | 11 (37.9%) |
| Mendoza | 134 (18.8%) | 3 (10.3%) |
| Neuquén | 112 (15.7%) | 9 (31.0%) |
| Santa Fe | 81 (11.3%) | 1 (3.5%) |
| La Pampa | 57 (8.0%) | 3 (10.3%) |
| Buenos Aires City | 36 (5.0%) | 1 (3.5%) |
| Rio Negro | 3 (0.4%) | 1 (3.5%) |
| Cases received from referral units, n (%)<sup>b</sup> | 275 (38.5%) | 17 (58.6%) |
| Cases received from referral units per<br>geographic location, n (%)<sup>c</sup> |  |  |
| Buenos Aires Province | 117 (40.2%) | 6 (54.6%) |
| Mendoza | 43 (32.1%) | 1 (33.3%) |
| Neuquén | 59 (52.7%) | 7 (77.8%) |
| Santa Fe | 34 (41.9%) | 1 (100%) |
| La Pampa | 19 (33.3%) | 1 (33.3%) |
| Rio Negro | 3 (100%) | 1 (100%) |
| Cases that received antibiotic treatment<br>before hospital admission, n (%)<sup>b</sup> | 30 (4.2%) | 0 (0.0%) |

<sup>a</sup>Abbreviations: IQR, interquartile range; SD, standard deviation; BD, bloody diarrhea; STEC: Shiga toxin-producing *Escherichia coli*
<sup>b</sup>Percentage is relative to the total number of cases reported within each group (Total BD or STEC-positive BD)
<sup>c</sup>Percentage is relative to the total number of cases reported at each Province and within each group (Total BD or STEC-positive BD)

**TABLE 2.**
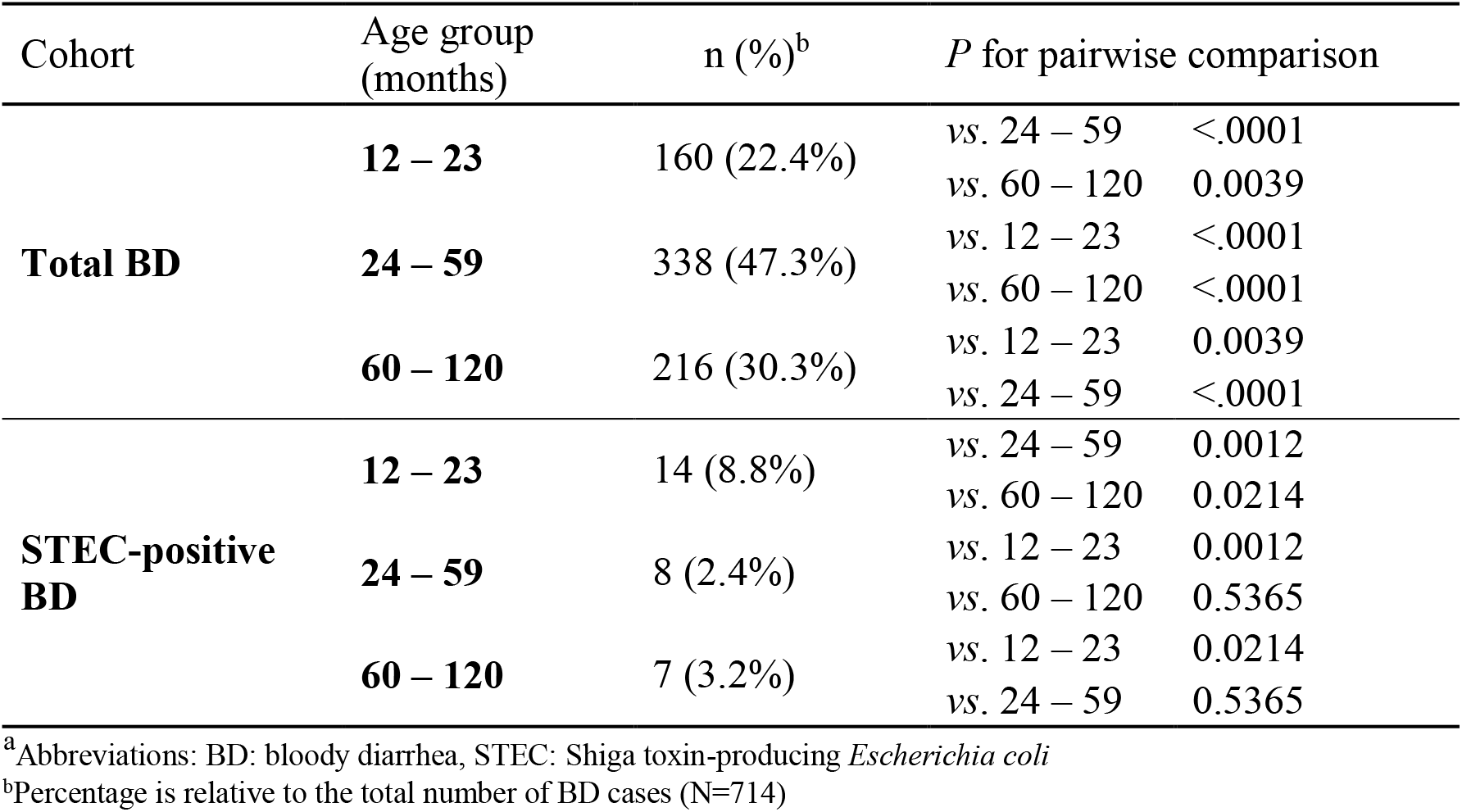
Distribution of BD and STEC-positive BD cases by age group^a^

| Cohort | Age group (months) | n (%)<sup>b</sup> | *P* for pairwise comparison |  |
| --- | --- | --- | --- | --- |
| **Total BD** | **12 – 23** | 160 (22.4%) | vs. 24 – 59 | <.0001 |
|  |  |  | vs. 60 – 120 | 0.0039 |
|  | **24 – 59** | 338 (47.3%) | vs. 12 – 23 | <.0001 |
|  |  |  | vs. 60 – 120 | <.0001 |
|  | **60 – 120** | 216 (30.3%) | vs. 12 – 23 | 0.0039 |
|  |  |  | vs. 24 – 59 | <.0001 |
| **STEC-positive BD** | **12 – 23** | 14 (8.8%) | vs. 24 – 59 | 0.0012 |
|  |  |  | vs. 60 – 120 | 0.0214 |
|  | **24 – 59** | 8 (2.4%) | vs. 12 – 23 | 0.0012 |
|  |  |  | vs. 60 – 120 | 0.5365 |
|  | **60 – 120** | 7 (3.2%) | vs. 12 – 23 | 0.0214 |
|  |  |  | vs. 24 – 59 | 0.5365 |

<sup>a</sup> Abbreviations: BD: bloody diarrhea, STEC: Shiga toxin-producing *Escherichia coli*
<sup>b</sup> Percentage is relative to the total number of BD cases (N=714)

The number and percentage of STEC-positive BD in each site were: Hospital “Dr. Castro Rendón” (n=10, 8.7%), Hospital “Dr. José Penna” (n=7, 7.9%), Hospital “Dr. Lucio Molas” (n=3, 5.3%), Hospital Italiano (n=3, 3.6%), Hospital “Dr. Humberto J. Notti” (n=3, 2.2%), Hospital “Sor María Ludovica” (n=2, 1.3%) and Hospital “Dr. Orlando Alassia” (n=1, 1.3%). Of the 29 STEC-positive BD detected, 17 (58.6%) were attended at the Referral Units and 12 (41.4%) at the Reference Sites (Fig. 1 and 2).

**FIG 2.**
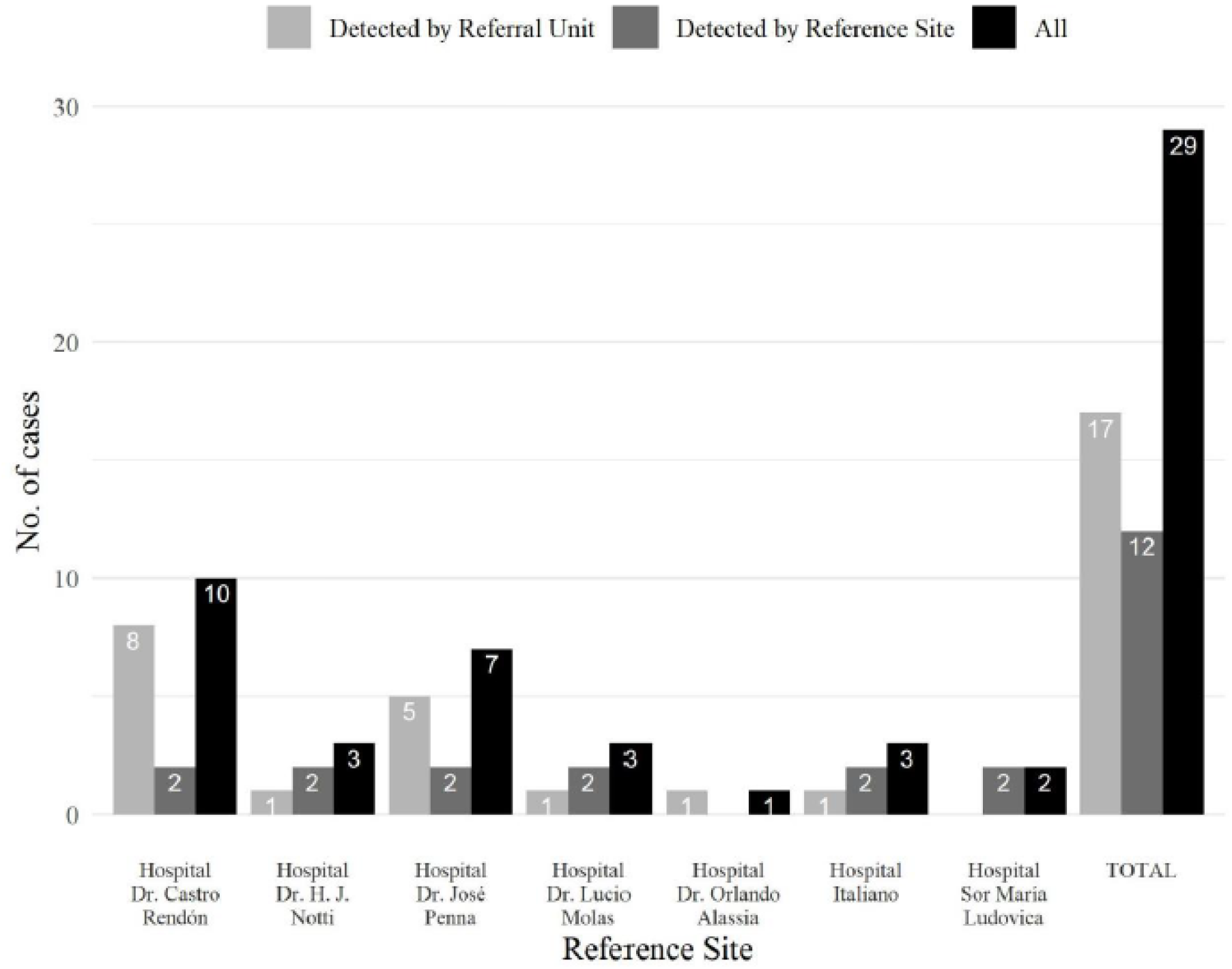
Frequency of STEC-positive bloody diarrhea cases by Reference Site

The proportion of STEC-positive BD over total BD cases by month was October, 0 (0.0%); November, 1 (4.2%); December, 2 (3.6%); January, 4 (2.8%); February, 9 (5.3%); March, 5 (4.6%); April, 3 (3.8%); May, 3 (3.9%) and June, 2 (3.6%). The highest frequencies occurred during February and March.

Among the 29 STEC-positive BD cases, 9 (31.0%) were O157:H7, and 6 (20.7%) were O145:H28. Other serotypes detected were O26:H11 (4), O121:H19 (1), ONT:H2 (1), ONT:HNT (3). In 5 cases, the *stx*-isolates could not be recovered and serotyping/genotyping could not be performed. Among the 24 STEC isolates, the frequency of the *stx*-genotypes was, *stx*_2a_ (n=12, 50.0%), *stx*_2a_/*stx*_2c_ (n= 8, 33.3%), *stx*_1a_ (n= 2, 8.3%), *stx*_1a_/*stx*_2a_ and *stx*_1a_/*stx*_2c_ (n= 1, 4.2%, each one). All O157:H7 strains were recovered during warm months.

Three (10.7%) STEC-positive BD cases, diagnosed by direct or enriched STQC and confirmed by mPCR at the local laboratories from Buenos Aires, Neuquén and Santa Fe developed HUS during follow-up. Additionally, one STEC-negative BD case by direct and enriched STQC tests and mPCR at the local laboratory in Mendoza, developed HUS 4 days after admission and STEC O157:H7 *stx*_2a_/*stx*_2c_, and anti-O157-specific antibodies were detected at the National Reference Laboratory. Thus, with the addition of this latter case, 13.8% of STEC-positive BD patients developed HUS, of whom three were ≤ 24 months of age. Progression to HUS occurred between three to five days after the BD onset during warm months. The median interval between the BD onset and the progression to STEC-HUS was 4.5 days (IQR 3-5 days; range, 3-5 days). STEC O157:H7 *stx*_2a_/*stx*_2c_ was isolated from three patients, while an STEC ONT:H2 *stx*_1a_/*stx*_2a_ was identified in the remaining one.

The 27 HUS patients of the second cohort were attended in the following hospitals: Sor María Ludovica (13), Notti (6), Penna (4), Castro Rendón (2), Italiano (1), Molas (1). The distribution by month was the following: December (3), January (8), February (9), March (5), May (1) and June (1).

The patient attended at the Hospital Italiano of Buenos Aires City was STEC-positive at the local laboratory of the Hospital of Trelew in Chubut Province, where the patient was first attended. Of the remaining 26 cases analyzed by STQC and mPCR, 11 (44%) were Stx-positive by STQC, 5 by direct method, and other 6 after enrichment method, while all 26 cases were positive by mPCR. None of the HUS patients received antibiotic treatment.

Among the 27 patients, STEC O157:H7 strains were recovered in 11 (40.7%), O145:H28 in 5 (18.5%). The remaining isolates were non-typeable. Among the O157:H7 isolates, the *stx*_2a_/*stx*_2c_ genotype prevailed (n=8) and all O145:H28 strains were *stx*_2a_-only. Two HUS cases were *stx*_2_-PCR positive without isolation. FFStx was detected in 7 out of 21 cases analyzed; anti-O157 antibodies were identified in 8 and anti-O145 in 5 out of 18 cases studied. The remaining 5 antisera samples were negative for the currently available antibodies (O157, O145, O121).

## DISCUSSION

Considering the impact of STEC-HUS on Public Health in Argentina, this study was carried out in order to assess the incidence of STEC-associated acute bloody diarrhea in children aged 1 to 9 years old, attended in seven tertiary-care hospitals and 18 Referral Units of different regions of the country.

Previous studies conducted in our country at various sites by López et al. (21) and Rivero et al. (22) reported a frequency of 4.1% and 10.06% STEC infections, respectively, in children with watery or bloody diarrhea. The health care centers participating in those studies were mainly located in the Central region of Argentina and only the first study reported the STEC incidence in the BD group, which was 9.6% (21). In comparison, we found STEC in a lower proportion of cases overall, 4.1% of the samples from patients with bloody diarrhea, but the incidence varied greatly among the different sites, ranging from 1.3% in La Plata and Santa Fe to 7.9% and 8.7% in Bahía Blanca and Neuquén. The latter two sites are located in the Southern region, with recognized high rate of HUS (https://www.argentina.gob.ar/sites/default/files/biv_463_cuatrimestral.pdf). However, the higher incidence of STEC-positive BD found could also respond to the fact that these two sites worked together with numerous Referral Units, where patients tend to seek for primary medical attention. Often, these patients arrive to tertiary hospitals with HUS already installed, and in this settings BD is less often seen.

Regarding age, the highest proportion of STEC-positive BD cases was found among patients of 12-23 months old, compared to the other two, from 24 to 60 and from 60 to 120 months. This finding is in agreement with other studies conducted in Argentina, which showed children under two years old as the most affected among patients with STEC infections (21, 22). Although there was a slightly larger proportion of males in the population studied, the gender distribution in the STEC-positive BD cases was even.

Antibiotic administration to individuals with STEC infections remains controversial, as stated in the meta-analysis by Freedman et al. (23), however the authors concluded that the use of such drugs is not recommended. In the present study, 4.2% of 714 patients with BD were prescribed with antibiotics, but none of the patients presenting STEC infections, either BD or HUS, received antibiotic treatment.

Most of the STEC-positive BD cases, including all *E. coli* O157:H7 cases, as well as the majority of HUS cases occurred during the summer months; this is in line with previous findings about seasonality, both in Argentina and other countries like USA (12, 13, 24, 25).

It is estimated that around 10% to 15% of patients develop HUS following STEC infection in the 2 weeks after the diarrhea onset, depending on the region and the season, and it is known that early recognition of infected patients is critical to improve the overall patient outcome and to be able to respond to outbreak situations in an effective and timely manner (26, 27).

In the present study, four (13.8%) STEC-positive BD patients developed HUS three to five days after the BD onset, during summertime. This rate is similar to those found in previous studies (28, 29). Three of these patients were aged ≤ 24 months. None received antibiotic treatment during the prodromal stage. The fecal samples were taken the same day of the bloody onset or one day later and processed by direct or enriched STQC and a validated in-house mPCR. All STEC isolated were Stx2a-producing strains in combination with other *stx* subtypes and belonged to O157:H7 (3) and ONT:H2 (1) serotypes. These results are in agreement with previous studies, in which *E. coli* O157:H7 harboring the *stx*_2a_/*stx*_2c_ genes was prevalent among diarrhea and HUS pediatric patients in Argentina (19, 30, 31). Recently, Adams et al. (32) published a study of 1,059 pediatric STEC patients seen in the UK between October 1, 2011, and October 31, 2014, of whom 207 developed HUS. They found that the predictors of risk of progression to HUS varied with age, *stx* type, antibiotic exposure, and clinical presentation, being children aged 1-4 years infected with *stx*_2_-only, with reported antibiotic exposure and presenting with bloody diarrhea or vomiting at highest risk. Tarr et al. (33) found in 936 *E. coli* O157:H7 cases a substantial increase in HUS risk associated with the *stx*_2a_ genotype as compared to the *stx*_1a_/*stx*_2a_ genotype, two of the most common genotypes in USA and Japan (34, 35, 36). Tarr et al. (33) also found that the virulence of the *stx*_2a_/*stx*_2c_ genotype, predominant in other settings (37, 38), was similar to *stx*_2a_-only genotype.

In Argentina, STEC-HUS is one of the most common etiologies of acute kidney injury in children, mortality rates can reach up to 3.6% (39), and long-term renal sequelae remains, on average, around 30-50% (40). For this reason, early and accurate diagnosis of STEC infection would be beneficial for early initiation of supportive treatment and the majority of STEC infected subjects tend to arrive at medical facilities already exhibiting bloody diarrhea. Moreover, it is known that the isolation rate of STEC in feces declines quickly after the first symptoms (41). In this study, only 40.5% of stool samples from patients with BD were collected on the same day of the BD onset, pointing out the need to strengthen public awareness regarding early consultation, as well as to improve the health system response so that both pediatricians and laboratory staff make sure to order and process sample collection timely.

Of the diagnostic techniques applied, STQC is a rapid membrane EIA that allows detection of the most common Stx1 and Stx2 subtypes, easy to perform and interpret. Chui et al. (2015) (42) reported a sensitivity and specificity of 70.0% and 99.4%, respectively, on 784 stool samples collected in southern Alberta, Canada and found a positivity rate of STEC infection of 2.6%. When enriched culture was used, the sensitivity increased to 85.0%. In our study, the STQC showed lower sensitivity for clinical specimens comparing with the results obtained using assays based on nucleic acid amplification as the in-house validated mPCR both for the samples from patients with BD as well as from those diagnosed with HUS. The use of an enrichment step improved the STQC performance, raising from 44.8% STEC-positive BD cases identified by the direct method to 72.4%, in line with previous reports (42, 43).

Among the 27 STEC-HUS patients of the second cohort, 77.8% were under 5 years old and 51.9% female, like those reported for Argentina (https://www.argentina.gob.ar/sites/default/files/biv_463_cuatrimestral.pdf). Regarding the origin, 63% resided in different locations of Buenos Aires province, 22.2% in Mendoza province, and the rest in Chubut, La Pampa, Neuquén and Río Negro provinces (3.7% each). In 26 HUS patients, 11 (44%) were Stx-positive by STQC, 5 by direct method, and other 6 after enrichment method, while all 26 cases were positive by mPCR. Historically in Argentina, the evidence of STEC infection in HUS patients was around 30%, but such diagnostic performance had significantly improved since 2014, reached >60% with the incorporation of anti-O serogroup-specific antibodies O157, O145, O121 detection (20). In the present study, anti-O157 antibodies were identified in 8 and anti-O145 in 5 of the serum samples available from 18 patients.

The serotypes and genotypes of the STEC strains isolated in the present study are in agreement with previous reports in Argentina. During a prospective study performed in Buenos Aires City and Mendoza, Rivas et al. (19) included 99 children with diarrhea and HUS. The 103 STEC strains isolated belonged to 18 different serotypes, and O157:H7 was the most frequent (59%), followed by O145:NM (12.6%), O26:H11 (5.8%), among others. Stx2 was identified in 90.3%, and Stx1 in 9.7% of the strains. Among the 61 STEC O157 strains, 93.4% harbored the *stx*_2a_/*stx*_2c_ genotype. More recently, Pianciola et al. (30) studied 280 O157 strains (54 bovine and 226 human) isolated between 2006 and 2008 in different regions of Argentina and found the *stx*_2a_/*stx*_2c_ genotype as prevalent in human (76.1%) and bovine (55.5%) strains. Moreover, 87.6% of the strains belonged to the hypervirulent clade 8. These data may help to understand the causes of the epidemiological situation related to HUS in Argentina.

In summary, 4.1% of BD cases was associated with STEC infections, with a great variability among the different regions of Argentina. Moreover, 13.8% of these STEC-positive BD cases evolved to HUS. Considering the endemic behavior of HUS in our country and its impact on public health, it is important that professionals have updated information about the epidemiology of the diarrheal disease, so that an accurate diagnosis can be made in a timely manner and patients have the best opportunity for a positive outcome.

## Acknowledgements

The authors thank all the professionals, technicians and assistants who collaborated in this multicenter study.

This work was supported by funds from INMUNOVA S.A.

